# Serotype specific anti-capsular polysaccharide IgG associated with colonizing and invasive group B streptococcus in pregnant woman-newborn dyads

**DOI:** 10.64898/2026.11.02.763839

**Authors:** Narjis Amar, Tom Hall, Emily Bentley, Azeddine Chakroun, Hanane Ben-Taleb, Rachid Saïle, Kirsty Le Doare, Mustapha Chamekh

**Affiliations:** Laboratory of Pediatric Research, Inflammation Unit, Faculty of Medicine, and ULB center for immunology, Université libre de Bruxelles (ULB), Belgium; Laboratory of Biology and Health, Research Center of Biotechnology and Health, Faculty of Sciences Ben M’Sik, Hassan II University of Casablanca, Morocco; Centre for Neonatal and Paediatric Infection, City St George’s University of London, London, United Kingdom

**Author notes:** Corresponding author Mailing address: Building V, Avenue Jean Joseph Crocq, 1-3, 1020 Brussels, Belgium.

**Keywords:** Group B Streptococcus, Anti-capsular polysaccharide IgG, Colonization, Early onset disease

## Abstract

Maternal IgG against group B streptococcus (GBS) capsular polysaccharide (CPS), is the principal placental mediator of neonatal protection, yet naturally acquired anti-CPS immunity has not been characterized in North African populations. To address this gap, we conducted a prospective study in a Moroccan mother-newborn cohort to quantify the serotype-specific anti-CPS IgG levels in relation to colonizing versus invasive GBS serotypes. Additionally, we assessed the transplacental transfer of anti-CPS IgG and identified potential correlates of invasive disease.

We examined serotype-specific anti-CPS IgG (serotypes Ia-V) in the plasma from maternal and cord blood of 103 GBS-colonized pregnant women, 35 uncolonized controls and their paired newborns (90 paired colonized dyads, including five newborns with invasive GBS disease) using a standardized six-plex immunoassay. Transfer ratios were compared with 1.0 and cord concentrations with published serotype-and onset-specific risk-reduction thresholds.

Maternal anti-CPS IgG concentrations varied by serotype, with the highest median for serotype II(1.36µg/mL) and the lowest for serotype Ib(0.003µg/mL). They were higher in GBS-colonized than uncolonized mothers for serotypes Ia, Ib and II. Maternal and cord blood IgG concentrations were correlated across serotypes, but placental transfer was incomplete, with median ratios of 0.50-0.80. Among 77 newborns, 13.0%(n=10) reached early-onset and 44.2%(n=34) achieved late-onset disease risk-reduction thresholds. The serotype IV was the most identified among neonatal infection disease cases, and this was associated with sub-protective anti-serotype IV antibody. This suggests that while naturally acquired anti-serotype II antibodies were predominant, the low level of anti-serotype IV antibodies could contribute to neonatal infection diseases in mother-newborn pairs.

## Introduction

Maternal immunization has become one of the most effective strategies for protecting the newborn during the first weeks of life, when the infant’s adaptive immune system cannot yet mount a timely response to invasive pathogens. Antigen-specific immunoglobulin G (IgG) generated in the mother is actively transported across the placenta through the neonatal Fc receptor on the syncytiotrophoblast accumulating in the fetal circulation from the early second trimester through the third [1,2]. GBS is regarded as one of the leading targets for maternal vaccine development [3,4].

GBS remains a major cause of neonatal sepsis, pneumonia and meningitis, with the burden concentrated in low- and middle-income regions where intrapartum antibiotic prophylaxis is difficult to deliver reliably and cannot prevent late-onset disease [5]. The principal mediator of protection is maternal IgG directed against the GBS capsular polysaccharide (CPS), a relationship first established by Baker and Kasper [6] and subsequently refined by sero-epidemiological studies linking higher maternal or infant anti-CPS IgG concentrations to a reduced risk of invasive disease, using the same standardized GASTON (Group B Streptococcal Assay Standardization) multiplex immunoassay (MIA), allowing direct comparison of concentrations across cohorts [7–14]. Naturally acquired anti-CPS responses are serotype-specific and functionally heterogeneous [15], and placental transfer is further reduced in preterm and pathological pregnancies, widening the gap between maternal concentrations and neonatal protection [2].

Both GBS colonization prevalence and serotype distribution vary markedly by region [19], highlighting the need to define GBS epidemiology in North African populations. Equally, naturally acquired maternal anti-CPS IgG concentrations and their placental transfer remain poorly characterized in this region, limiting our understanding of naturally acquired protection in newborns. Recently, in a cohort of pregnant women we reported a GBS carriage prevalence of 9.9%, with serotypes Ib (19.8%), IV (17.1%), II (14.3%), V (13.7%), Ia (12.3%) and III (7.5%) in decreasing order of frequency [17]. Furthermore, we showed that GBS-colonized-mothers exhibit higher levels of inflammatory cytokine response compared to GBS uncolonized-mothers [20]. Interestingly, maternal IL-17A, a crucial mediator of antibacterial defense, emerges as a significant prognostic biomarker for evaluating the risk of GBS transmission to neonates [20]. In this study, we aimed to quantify serotype specific anti-CPS IgG concentrations in mother-newborn dyads from the same cohort using the GASTON MIA. We also assessed the placental transfer of these antibodies and their association with neonatal invasive GBS disease.

### Methods Study design

The present work is a serological sub-study conducted within a prospective mother-newborn cohort previously described by Ben-Taleb and colleagues [17], who reported the capsular serotyping data, and by Amar and colleagues [20], who established maternal IL-17A as a prognostic biomarker for invasive neonatal GBS disease. The present sub-study quantifies capsular polysaccharide-specific IgG (anti-CPS IgG) in the same maternal and neonatal cord plasma samples.

A total of 270 plasma samples were analyzed drawn from four groups: GBS-colonized pregnant women, newborns with invasive GBS disease, healthy newborns born to colonized or uncolonized mothers, and uninfected controls. Participants were recruited at the maternity unit of the Abderrahim Harouchi Mother-Child Hospital, Averroes University Hospital Center, Casablanca, Morocco, between April 2021 and July 2024.

The study was approved by the local hospital ethics committee (N° 21/2021) and the National Commission for the Control of Personal Data Protection (N° A-RS-248/2021). Written informed consent was obtained from all participating women on their own behalf and on behalf of their newborns.

### Study population and clinical definitions

Maternal GBS screening was performed as described in Ben-Taleb et al. [17]. Briefly, vaginal swabs collected at 35-40 weeks of gestation, in line with CDC 2010 guidelines [21], were plated on Columbia agar with 5% sheep blood, nalidixic acid (15 µg/mL) and colistin (10 µg/mL) (bioMérieux, Marcy-l’Étoile, France) and incubated for 18-24 h at 37 °C. β-hemolytic colonies were confirmed biochemically (API® 20 Strep, bioMérieux) and identified as Lancefield group B by latex agglutination (Streptococcal Grouping Kit, Thermo Scientific™ Oxoid™, Basingstoke, UK).

Symptomatic GBS-colonized mothers were defined following the parent study [20] as GBS-positive culture with at least two inflammatory signs (intrapartum fever > 38 °C, elevated CRP, abnormal leukocyte count). For the integrated transmission analysis, colonized mothers were further grouped by neonatal outcome as non-transmitting (NTM; newborn with no clinical or biological signs of infection), suspected transmission (M-SI; newborn with at least two clinical or biological signs of infection (abnormal leukocyte count, elevated CRP and PCT) but a negative GBS blood culture), or transmitting (TM; newborn with blood-culture-confirmed GBS bacteremia (invasive disease). Uncolonized mothers whose newborns showed no signs of infection served as controls. A total of 90 maternal and 92 paired cord samples from colonized dyads, and 23 maternal and 24 paired cord samples from controls, were tested. For the composite analysis, symptomatic colonized women carrying a serotype included in the assay panel (n=33) were classified according to newborn outcome as NTM (n=3), M-SI (n=25), or TM (n=5). Inclusion and exclusion criteria followed the parent cohort [20].

### Sub-study sample selection

Of the 124 paired GBS-colonized mother-newborn dyads in the parent cohort [20], anti-CPS IgG concentrations were quantified in plasma from 103 colonized mothers. The remaining dyads were excluded because insufficient residual plasma was available following prior ELISA and Luminex analyses. 90 of these pregnant women had paired cord-blood samples and form the analytic paired cohort, yielding 92 newborns (one triplet pregnancy); the remaining 13 contributed a maternal sample only. Of 35 uncolonized control women, 23 had a paired cord-blood sample, yielding 24 newborns (one twin pregnancy); 12 contributed a maternal sample only. Uncolonized controls were randomly selected from 150 GBS-negative women who delivered in the study unit and for whom paired maternal and cord samples were available, as described from the parent cohort [20]. Among the 90 colonized women with paired maternal and cord samples, 75 carried one of the six serotypes measured by the GASTON MIA (Ia (n=10), Ib (n=18), II (n=17), III (n=6), IV (n=15), V (n=9)); five carried serotype VII, two serotype VI, two were non-typeable and six were not typed. Five newborns developed culture-confirmed invasive GBS disease, identified by positive neonatal blood culture, and all five were early-onset. Isolates were capsular typed in three of five cases and all three were serotypes IV; for the two untyped cases the maternal colonizing serotype (V and Ia) was used as the homologous reference.

### Sample collection and processing

Maternal peripheral blood was drawn on admission, 5-7 hours before delivery, and cord blood was collected at delivery. Samples were centrifuged at 200 × g for 10 min at 4 °C within 4 h of collection; plasma was aliquoted and stored at -80 °C until analysis.

### Cytokine concentrations (data re-used from the parent study)

Plasma concentrations of IL-1β, IL-17A and IL-4 in maternal and cord blood were not re-measured in the present sub-study. The values used in this study were previously generated and reported for the parent cohort [20], in which the protocol for cytokine quantification in maternal and cord-blood plasma collected at delivery is described in detail. Here, cytokine data are used as a secondary readout of disease severity across outcome strata; the primary analyses concern the anti-CPS IgG data generated in this sub-study.

Concentrations below the serotype-specific lower limit of quantification (LLOQ; Ia, 0.002; Ib, 0.005; II, 0.022; III, 0.009; IV, 0.004; V, 0.010 µg/mL) were assigned half the LLOQ (Ia, 0.001; Ib, 0.0025; II, 0.011; III, 0.0045; IV, 0.002; V, 0.005 µg/mL). A sensitivity analysis in which values below the LLOQ were substituted with either the full LLOQ or zero did not alter any of the threshold-based results. Dyads in which the maternal concentration was below the serotype-specific LLOQ were excluded from the transfer-ratio analysis, as transfer ratios could not be reliably estimated when the maternal concentration used as the denominator fell below the quantifiable range of the assay.

### Capsular serotyping of GBS isolates

GBS isolates from maternal vaginal swabs and positive neonatal blood cultures were serotyped by multiplex PCR using the primers of Poyart and colleagues [22], with *recA* or 16S rRNA as internal amplification controls, as described in Ben-Taleb et al. [17]. Genomic DNA was extracted from fresh cultures using the Canvax HigherPurity™ Bacterial Genomic DNA Isolation Kit (Canvax Biotech S.L., Cordoba, Spain) and stored at -20 °C until amplification. Amplicons were resolved by agarose gel electrophoresis. Isolates were assigned to serotypes Ia, Ib, II, III, IV, V, VI or VII, or classified as non-typeable (NT) when no serotype-specific amplicon was observed despite a positive internal control.

### Quantification of anti-CPS IgG: the GASTON multiplex immunoassay

Serotype-specific IgG against the six GBS CPS serotypes responsible for the majority of invasive infant disease (Ia, Ib, II, III, IV and V) was measured using the GASTON assay, a six-plex Luminex-based MIA developed by Pfizer (Pearl River, NY, USA), validated for the quantification of anti-CPS IgG in human serum [23] and standardized across multiple labs [24]. In this study, the assay was applied to plasma samples. Samples from the Moroccan cohort were shipped to City St George’s, University of London (CSG; London, UK), where all sample testing was performed.

The GASTON MIA is based on Luminex MagPlex xMAP technology: six spectrally distinct, fluorescently dyed magnetic microspheres are each coupled to one of the six GBS CPS antigens via a poly-L-lysine linker, allowing simultaneous quantification of IgG against all six serotypes in a single well. The full assay procedure has been described in detail [23].

### Statistical analysis

Distributions were assessed by Shapiro-Wilk test on log₁₀-transformed concentrations. Transformation did not normalize the data for five of six serotypes (p<0.05), and therefore non-parametric methods were used throughout; concentrations are summarized as medians with interquartile ranges. Between-group comparisons were performed with the Mann-Whitney U or Kruskal-Wallis test, with Dunn’s post-hoc or Holm-Šídák correction for multiple groups, depending on distribution. Paired maternal and cord IgG concentrations were correlated using Spearman’s rank correlation coefficient. Transfer ratios were compared with the reference value of 1.0 using the Wilcoxon signed-rank test on log-transformed ratios. For the integrated analysis, a composite maternal protection score was defined for each mother as the sum of the z-transformed log₁₀ serotype-matched anti-CPS IgG and the z-transformed log₁₀ maternal IL-17A, with higher values indicating greater protection. Scores were compared across newborn-outcome groups by the Kruskal-Wallis test with Dunn’s post-hoc correction.

Cord concentrations were compared with published serotype-specific thresholds associated with an 80% reduction in the risk of invasive disease, reported separately for early- and late-onset disease [13]. Because all five invasive cases in this cohort were early-onset, early-onset thresholds were applied (Ia, 2.49; Ib, 1.13; II, 12.91; III, 0.52; IV, 2.85; V, 5.25 µg/mL). Thresholds were applied to cord rather than maternal concentrations, since the newborn is the population at risk. Categorical comparisons used the chi-square or Fisher’s exact test.

A two-sided *p*-value < 0.05 was considered statistically significant. Analyses were performed in GraphPad Prism version 11.0.0 for macOS.

### Placental transfer ratio

For each maternal-newborn pair, a serotype-specific transfer ratio was calculated as the cord-blood IgG concentration divided by the paired maternal IgG concentration at delivery. Dyads in which the maternal IgG concentration was at the lower limit of the assay’s quantifiable range (≤ 0.005 µg/mL) were excluded, because a near-zero denominator yields artifactually inflated ratios; cord values at the limit of quantification were retained, and no upper limit was applied to the ratio. A ratio below 1 indicates incomplete transfer.

## Results

### Cohort characteristics

Serotype-specific anti-CPS IgG antibody concentrations were quantified against the six clinically relevant capsular serotypes Ia, Ib, II, III, IV and V in 103 GBS-colonized pregnant women and 35 uncolonized controls, together with 92 newborns of colonized mothers and 24 control newborns. Ninety pairs of maternal and cord-blood IgG samples were used as the analytic paired cohort for the transfer and threshold analyses. Five newborns had culture-confirmed invasive GBS disease, while 26 newborns exhibited clinical and/or inflammatory symptoms suggestive of suspected infection. Colonized and uncolonized pregnancies were comparable for maternal age, gestational age, preterm rate and neonatal birthweight, but colonized women had significantly higher systemic inflammatory markers; C-reactive protein, white-cell count, neutrophils and monocytes, and more frequent intrapartum fever (>38°C). Consistent with the parent cohort, the distribution of GBS serotypes in colonized mothers showed the predominance of Ib, II, and IV, whereas serotypes Ia, V, III, VII, and VI were less frequent (Table 1).

**Table 1.** Cohort characteristics, biological markers and serotype distribution.

| Characteristic | GBS-colonized mothers<br>n = 103 | GBS-uncolonized mothers<br>n = 35 | p value |
|---|---|---|---|
| **Maternal age, years (mean ± SD)** | 29.9 ± 7.1 | 31.4 ± 7.5 | 0.36 |
| **Gestational age (mean ± SD)** | 37.7 ± 3.3 | 38.5 ± 1.3 | 0.41 |
| **Parity - median (IQR)** | 2 (1-3) | 2 (1-2) | **0.011** |
| **C-section delivery, n/N (%)** | 30/80 (37.5%) | 11/16 (68.7%) | **0.027** |
| **Intrapartum fever >38°C, n (%)** | 17 (17%) | 1 (3%) | **0.043** |
| **Maternal CRP (mg/L)** | 35.1 ± 58.9 | 7.93 ± 4.85 | **0.004** |
| **Maternal leukocytes (10<sup>3</sup>/µL)** | 13.18 ± 3.2 | 11.11 ± 3.66 | **0.021** |
| **Maternal neutrophils (10<sup>3</sup>/µL)** | 10.74 ± 3.74 | 6.35 ± 3.55 | **<0.0001** |
| **Maternal monocytes (10<sup>3</sup>/µL)** | 0.72 ± 0.28 | 0.59 ± 0.23 | **0.041** |
| **Serotype distribution, n/N (%)** |  |  |  |
| **Ia** | 11/89 (12.4) | NA | — |
| **Ib** | 18/89 (20.2) | NA | — |
| **II** | 18/89 (20.2) | NA | — |
| **III** | 6/89 (6.7) | NA | — |
| **IV** | 17/89 (19.1) | NA | — |
| **V** | 9/89 (10.1) | NA | — |
| **VI** | 2/89 (2.2) | NA | — |
| **VII** | 6/89 (6.7) | NA | — |
| **NT** | 2/89 (2.2) | NA | — |
|  | **Newborns of colonized mothers<br>n=92** | **Newborns of uncolonized mothers<br>n=24** |  |
| **Birthweight (g)** | 2795 ± 719.4 | 2450 ± 788.1 | 0.067 |
| **Male sex, n/N (%)** | 51/92 (55.4%) | 12/24 (50%) | 0.65 |
| **Blood culture-positive (invasive) (n)** | 5 | 0 | — |
| **Newborn CRP (mg/L)** | 21.63 ± 39.69 | 3.08 ± 2.73 | **0.0077** |
| **Newborn PCT (mg/mL)** | 0.26 ± 0.57 | NA | — |
| **Newborn leukocytes (10<sup>3</sup>/µL)** | 12.13 ± 4.77 | NA | — |
| **Newborn neutrophils (10<sup>3</sup>/µL)** | 6.75 ± 4.23 | NA | — |
| **Newborn monocytes (10<sup>3</sup>/µL)** | 1.83 ± 2.19 | NA | — |

*Delivery mode was available for 80/103 colonized and 16/35 control mothers; percentages are calculated on these denominators Normally distributed variables are mean ± SD; skewed variables are median (IQR); categorical variables Fisher's exact test. Maternal counts treat each mother once; newborn counts retain siblings from multiple births (one colonized mother had triplets, one control mother had twins). CRP, C-reactive protein; PCT, procalcitonin.*

### Maternal IgG anti-GBS capsular polysaccharide

As shown in figure 1A, maternal anti-CPS IgG concentrations differed markedly across the six serotypes (p<0.0001). Median IgG concentration was highest for serotype II (1.36 µg/mL, IQR 0.24-7.01) and 28- to 47-fold lower for serotypes Ia, III, IV and V (medians 0.049, 0.048, 0.037 and 0.029 µg/mL, respectively). Serotype Ib concentrations were lowest, with more than half of the women having concentrations below the LLOQ (median 0.003 µg/mL).

For each dyad, cord IgG against the mother’s colonizing serotype was compared with the published concentration associated with an 80% reduction in invasive disease risk for that serotype, reported separately for early-onset and late-onset disease [13]. Because all five invasive cases in this cohort were early-onset, the early-onset thresholds are the primary comparison, although both are reported. Among the 77 newborns whose mother carried a serotype covered by the assay, 10 (13.0%) reached the early-onset threshold, and 34 (44.2%) reached the lower late-onset threshold. Coverage against the early-onset threshold varied by serotype: 5 of 17 (29.4%) for serotype II, 2 of 10 (20.0%) for Ia, 1 of 8 (12.5%) for III, 1 of 15 (6.7%) for IV, 1 of 18 (5.6%) for Ib, and none of 9 for serotype V. Coverage against the late-onset threshold was higher across all serotypes: 15 of 17 (88.2%) for serotype II, 8 of 15 (53.3%) for IV, 3 of 8 (37.5%) for III, 3 of 10 (30.0%) for Ia, 5 of 18 (27.8%) for Ib, and none of 9 for serotype V (Figure 1B).

**Figure 1.**
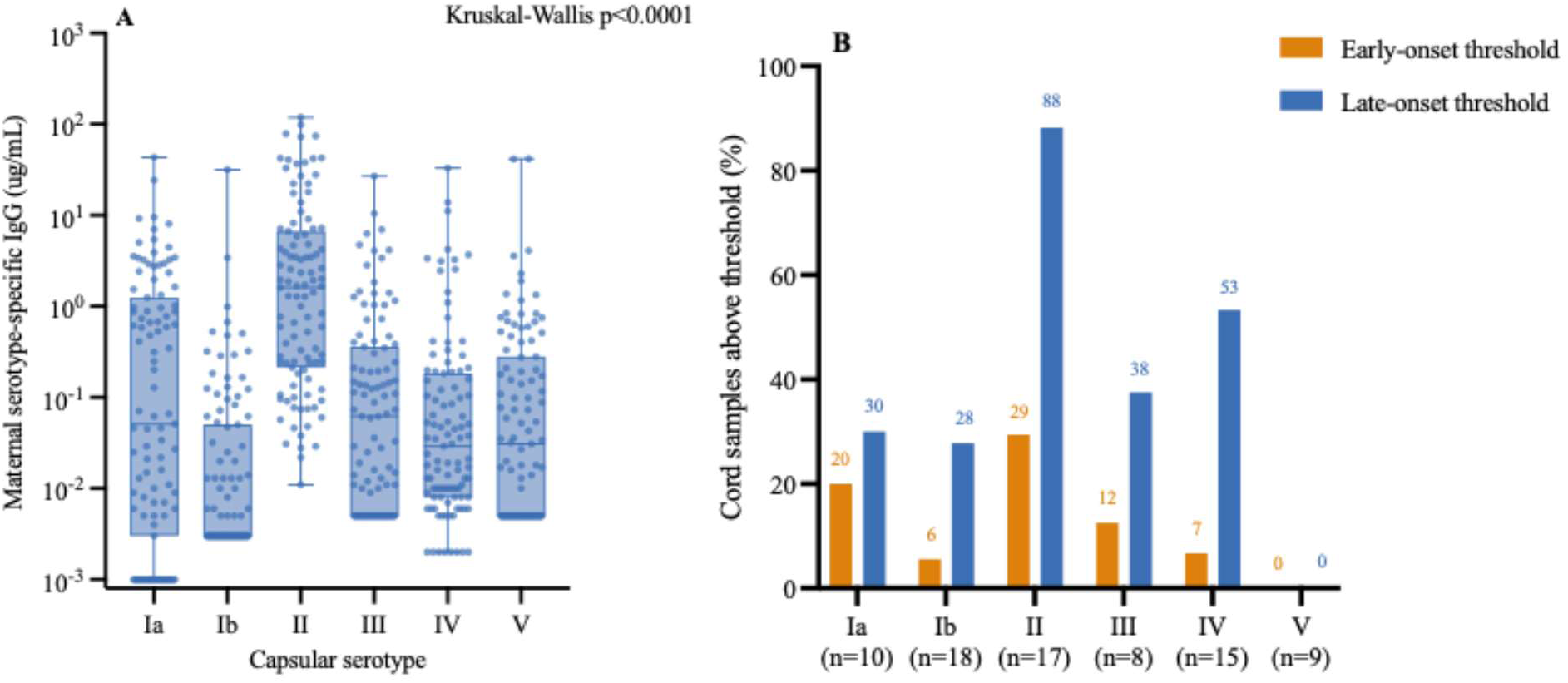
(A) Maternal serotype-specific anti-CPS IgG by capsular serotype (Kruskal-Wallis p<0.0001). Boxes show median and interquartile range. (B) Proportion of cord samples reaching the CDC early-onset (orange) and late-onset (blue) 80% risk-reduction thresholds [13], matched to the mother’s colonizing serotype. n = number of dyads per serotype

GBS-colonized mothers exhibited significantly higher level of IgG against serotypes Ia, Ib and II compared with GBS-uncolonized mothers (p<0.001, p<0.01 and p<0.02 respectively); while IgG anti-serotypes III, IV and V showed no significant difference despite the same trend. (Figure 2)

**Figure 2:**
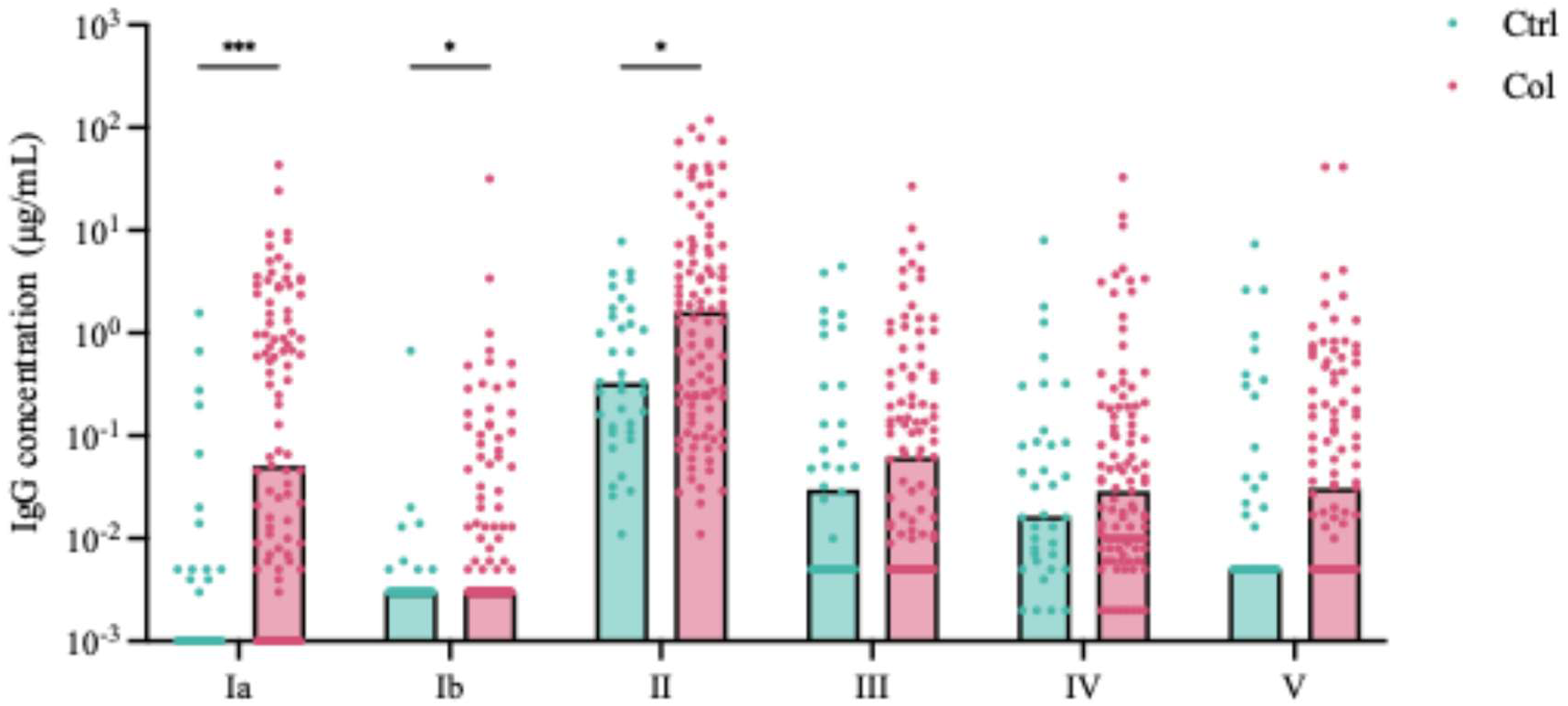
Distribution of serotype-specific anti-CPS IgG against for Ia, Ib, II, III, IV, and V in GBS-colonized (Col) and GBS-uncolonized mothers (Ctrl). Mann-Whitney test and Holm-Šídák correction for multiple comparisons. * p < 0.05, *** p < 0.001

Across all pregnant women, the homologous IgG response was significantly higher (median=0.14 µg/mL) than the heterologous IgG response (median = 0.035 µg/mL) (p=0.006) (Figure 3A).

The homologous anti-CPS IgG response varied markedly according to the maternal colonizing serotype (Figure 3B). Median homologous IgG concentrations were highest for serotype II (5.9 µg/mL; IQR 3.5-22.3; n=17), followed by serotype IV (0.33 µg/mL; IQR 0.03-2.9; n=15), V 0.11 (0.005-0.15, n=9), III 0.11 (0.03-0.30, n=6), Ia 0.01 (0.001-2.45, n=10), and Ib 0.008 (0.003-0.05, n=18). The homologous IgG response was significantly higher than the heterologous responses for serotypes II and IV (Mann-Whitney p=0.0003 and p=0.0104, respectively; Figure 3B), but not for the other serotypes.

**Figure 3.**
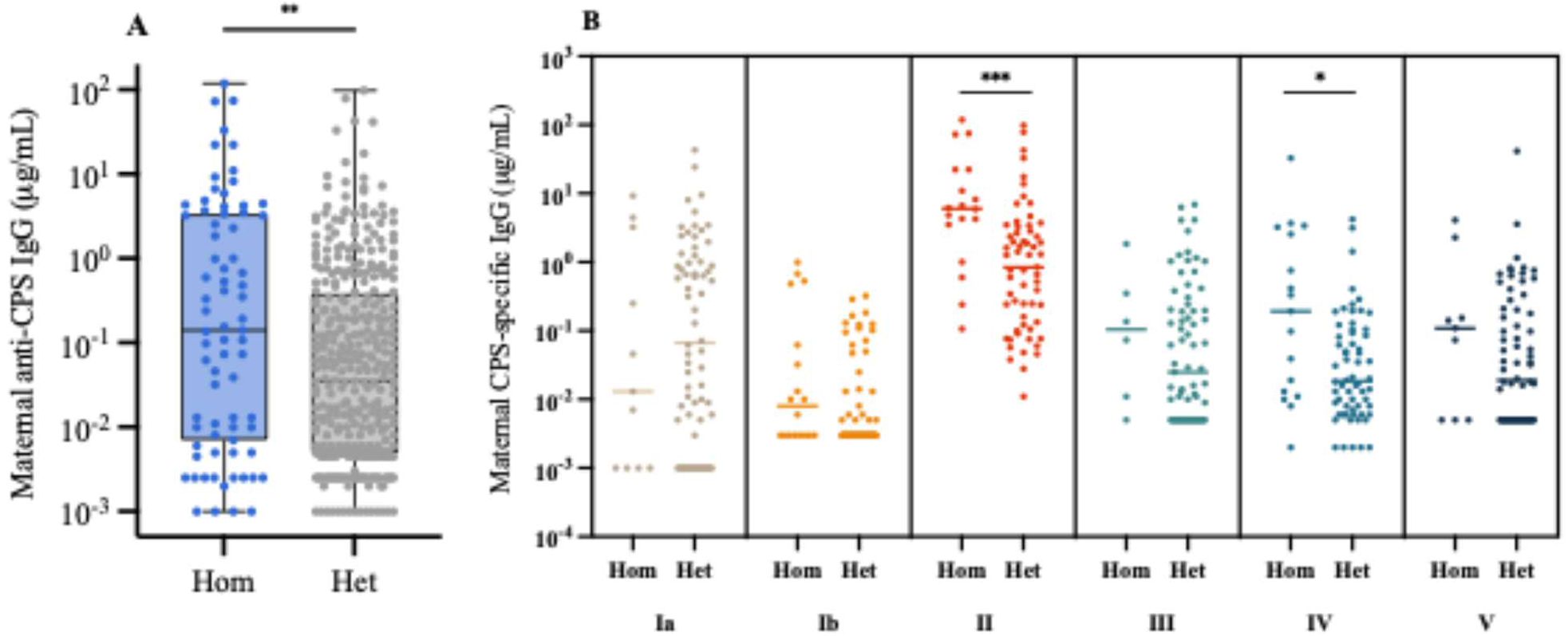
Homologous versus heterologous anti-CPS IgG response. (A) Maternal IgG against the colonizing serotype (homologous) versus other serotypes (heterologous). (B) Comparison of homologous versus heterologous Ig G response by colonizing serotype. Horizontal bars indicate median (± IQR). Het, heterologous; Hom, homologous. Mann-Whitney p shown. * P < 0.05, ** P < 0.01, *** P < 0.001.

### Cord-blood anti-GBS capsular polysaccharide IgG

Cord-blood IgG concentrations mirrored the maternal profile, with serotype II showing the highest median concentration (median 0.71 µg/mL), while concentrations for the other serotypes were generally low and close to the respective LLOQs (Figure 4A). Paired maternal and cord concentrations were strongly correlated for every serotype (Figure 4B-G; Spearman r=0.84-0.90, all p<0.0001; pooled r=0.91, 551 observations), confirming that neonatal anti-CPS IgG is overwhelmingly maternally acquired. Except for serotype II, cord IgG was generally lower than the paired maternal IgG, consistent with incomplete placental transfer.

**Figure 4.**
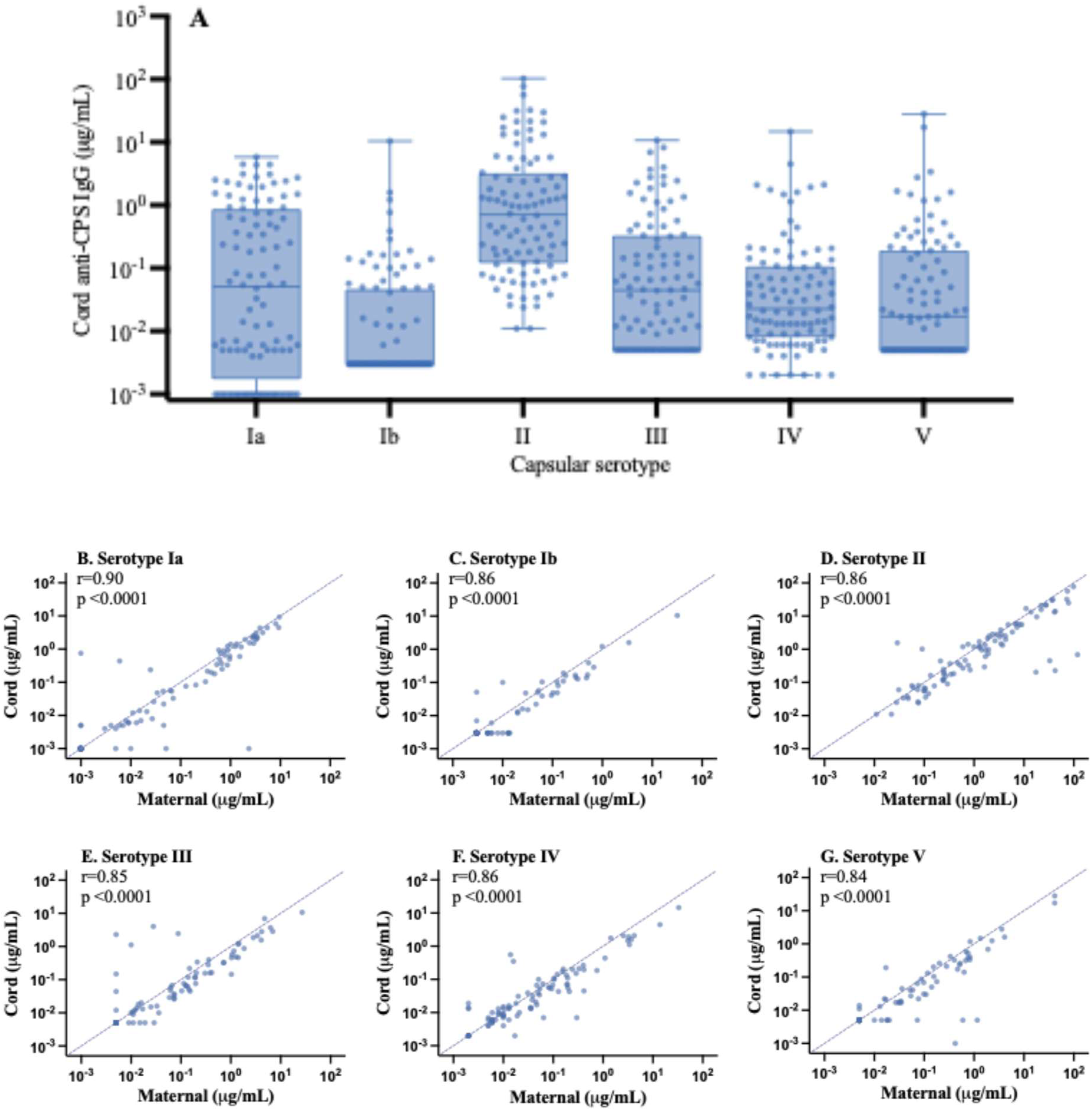
Cord-blood anti-CPS IgG and paired maternal-cord blood correlations (n=90). (A) Cord blood IgG against specific GBS serotypes. (B-G) Paired maternal versus cord blood IgG against specific GBS serotypes on log-log scales; dashed diagonal = equality. Spearman r per panel.

In contrast to the maternal IgG response, cord blood concentrations were similar for homologous and heterologous serotypes (median: 0.019 vs 0.019 µg/mL, p=0.40). The high homologous response against serotypes II and IV found in mothers was no longer apparent in cord blood, suggesting that the placental transfer was globally incomplete. Placental transfer ratios (cord/maternal IgG), recorded after excluding dyads with maternal IgG below the serotype-specific LLOQ, were below 1 for every serotype (medians: IV=0.80, Ia=0.71, II=0.70, III=0.58, V=0.53, Ib=0.50; Wilcoxon signed-rank vs 1.0, all p<0.001) (Figure 5).

Placental transfer did not differ by birthweight category (median ratio 0.65 versus 0.57, p=0.54) or delivery mode (vaginal 0.64 versus C-section 0.58, p=0.21), but correlated weakly with gestational age (Spearman ρ=0.21, p=0.04), in the expected direction of lower transfer at earlier gestations.

**Figure 5.**
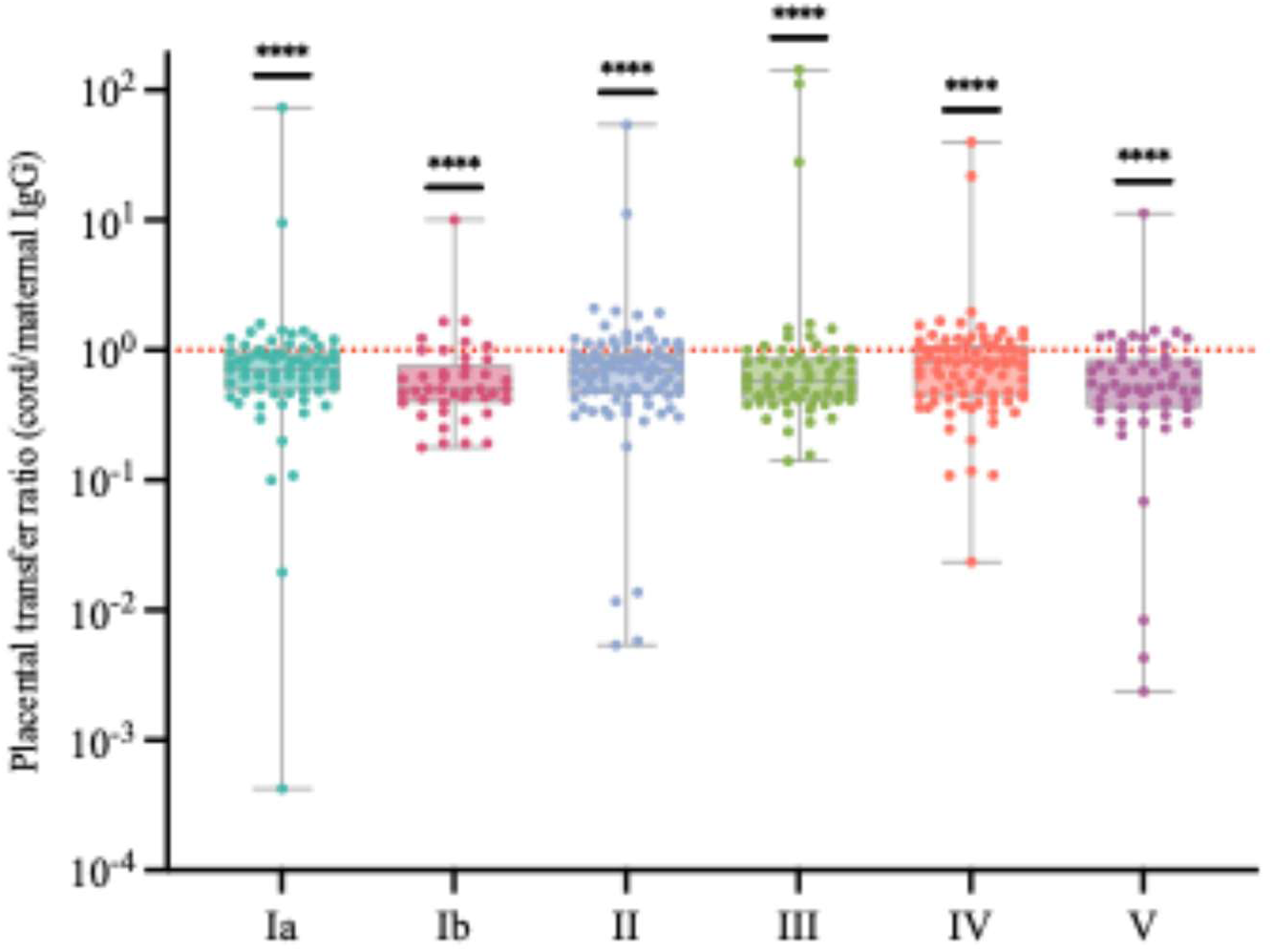
Placental transfer of anti-CPS IgG across different GBS serotypes. Transfer ratio (cord/maternal IgG) per serotype was recorded after excluding dyads with maternal IgG below the serotype-specific LLOQ. Boxes show median and IQR; the dashed line marks a ratio of 1.0 (equal cord and maternal concentrations). ****p<0.0001 versus 1.0 (Wilcoxon signed-rank test). n = 68 (Ia), 42 (Ib), 89 (II), 62 (III), 83 (IV), 55 (V).

### Invasive neonatal GBS disease is associated with deficiency of the maternal capsular antibody response

Five newborns developed culture-confirmed invasive GBS disease. Three isolates were serotype IV, matching the maternal colonizing serotype. Two could not be typed, due to a technical issue of the strain conservation. For these cases, maternal colonizing serotypes (Ia and V) were considered the likely homologous reference, as invasive disease typically originates from the strain the mother carries. Serotype IV was the most frequent serotype among the confirmed invasive cases. In all five invasive cases, homologous IgG concentrations were low in both maternal and cord blood (maternal median 0.008 µg/mL, range 0.002-0.331; cord median 0.006, range 0.002-0.211).

We have previously demonstrated in the parent cohort that lower maternal plasmatic level of IL-17 correlated with high risk of GBS neonatal invasive diseases [20]. We sought to analyze this parameter in conjunction with the level of anti-CPS IgG in GBS-colonized mothers who exhibited inflammatory symptoms and were at risk of transmitting the bacteria to their neonates. Among these symptomatic colonized women (n=33), both IL-17A and homologous anti-CPS IgG decreased with worsening outcome: NTM had the highest homologous IgG (median 8.2 µg/mL, IQR 4.2-15.3) and IL-17A (median 235 pg/mL, IQR 178-240); M-SI showed intermediate levels (IgG 0.24, IQR 0.01-4.11; IL-17A 76, IQR 43-120); and TM the lowest (IgG 0.008, IQR 0.005-0.013; IL-17A 38, IQR 34-85). A composite score [z(log₁₀ homologous IgG) + z(log₁₀ IL-17A)] decreased across NTM (median +2.87), M-SI (-0.26) and TM (-1.66) (Kruskal-Wallis p=0.013; Figure 6A). Given the small NTM group, this analysis is exploratory.

The two-dimensional distribution resulted in clear separate groups. Non-transmitting mothers were mainly at the antibody-replete, IL-17A-high region, while transmitting mothers were found at the antibody-deficient, IL-17A low region (Figure 6B). Invasive neonatal GBS disease occurred specifically where both maternal homologous response and IL-17A response failed simultaneously, rather than due to a deficiency in either response individually.

**Figure 6.**
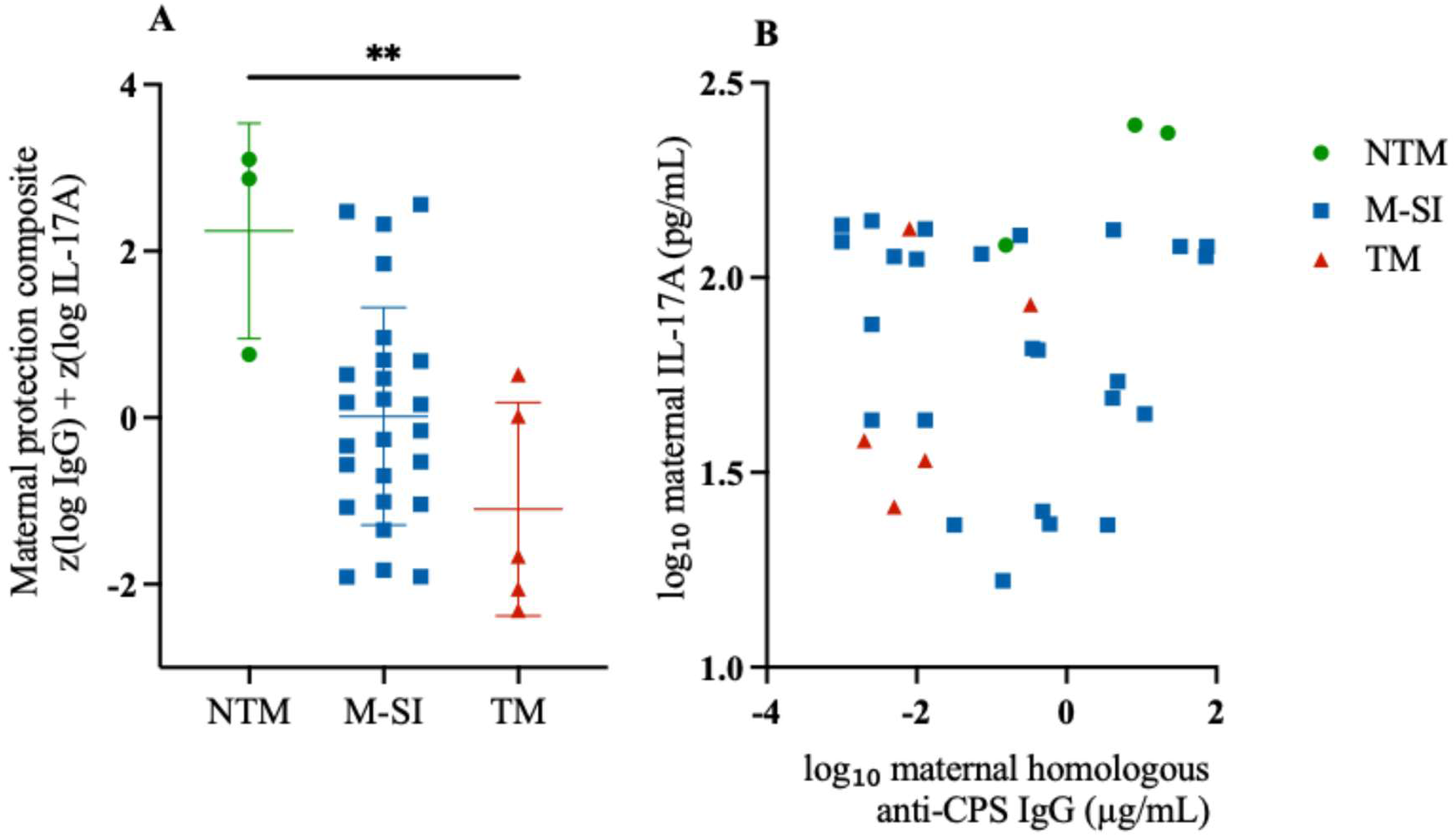
Invasive neonatal GBS disease coincides with a deficient maternal homologous anti-CPS IgG response. (A) Maternal protection composite [z (log₁₀ maternal homologous anti-CPS IgG) + z (log₁₀ maternal IL-17A)], related to newborn outcome: non-transmitting mothers (NTM, n=3), GBS-colonized-mothers with newborns with suspected infection (M-SI, n=25), and transmitting mothers (TM, n=5). Bars, median ± IQR; Kruskal-Wallis p=0.01, **p<0.01 NTM vs TM (Dunn’s correction). (B) Two-dimensional distribution: Each point is one mother, plotted by log₁₀ maternal homologous anti-CPS IgG versus log₁₀ maternal IL-17A.

## Discussion

The principal novel finding of this study is that a composite measure of maternal serotype-matched anti-CPS IgG and IL-17A distinguished newborn outcomes among symptomatic colonized women, with the lowest values observed in mothers whose newborns developed invasive disease. To our knowledge, this is the first paired mother-newborn analysis of naturally acquired anti-GBS capsular antibody in North Africa, and the first to combine humoral and mucosal markers in this way.

### Maternal anti-CPS IgG

Maternal anti-CPS IgG levels varied markedly between serotypes from a median of 1.36 µg/mL for serotype II to 0.003 µg/mL for serotype Ib. Anti-CPS IgG concentrations against serotypes Ia, Ib and II were significantly higher in GBS-colonized than in uncolonized women. Within colonized women, IgG concentrations against the carried serotype exceeded IgG concentrations against non-carried serotypes (0.14 versus 0.035 µg/mL, p=0.006), a serotype-specific pattern consistent with an antibody response elicited by maternal GBS carriage. In murine models of intestinal GBS acquisition, colonization alone elicited an endogenous serum anti-GBS IgG response within 20 days of exposure [38]. Our results are in line with previous European and African studies. In the large European DEVANI study, GBS-colonized women had higher IgG response against the carried serotype Ia, Ib, III, and V than non-carriers [14]. A UK study reported that anti-CPS IgG was 6.8-fold higher in GBS-colonized than uncolonized non-pregnant women [29]. The same pattern has been reported in African mother-newborn cohorts: in a Ghana-Zimbabwe cohort, homologous IgG for serotype Ia was significantly higher in GBS-colonized than uncolonized women in Ghana (0.61 versus 0.03 µg/mL, p=0.003) [30]. Higher serotype-specific IgG concentrations have likewise been reported in pregnant women colonized at delivery, although in that cohort the association held for serotypes Ia, II, III and V but not Ib [26]. In a longitudinal South African cohort, the relationship ran in the opposite direction: higher pre-existing serotype-specific IgG was associated with a lower risk of subsequently acquiring that same serotype [28]. The overall antibody response in our cohort was low for most serotypes, as reported in other African settings; in a South African case-control study, median maternal concentrations among colonized mothers of healthy infants were 0.29 µg/mL for both serotypes Ia and III, approximately six-fold lower than in North American controls [10]. Low natural antibody is not unique to low-income settings, naturally acquired anti-CPS IgG is also limited in high-income countries such as the UK [37]. These low absolute concentrations, seen here across most serotypes, explain the modest proportion of newborns reaching published risk-reduction thresholds [13] and support the rationale for maternal immunization.

Homologous IgG concentrations were highest for serotype II (median 5.9 µg/mL) and lowest for serotype Ib (0.008 µg/mL), despite similar carriage frequencies (n=17 and n=18), indicating that the magnitude of the homologous response does not simply reflect frequency of exposure. Serotype IV was notable in this cohort. It was among the most frequently carried serotypes in our earlier study of this cohort (17.1%) [17] and accounted for all three capsular-typed invasive isolates despite generally low homologous antibody concentrations. The homologous response exceeded heterologous responses significantly only for serotypes II and IV, but for serotype IV the difference was small in absolute terms and did not prevent invasive disease: the affected newborns were born to mothers with low serotype IV IgG (median maternal homologous concentration 0.008 µg/mL). A similar finding was reported in an earlier Canadian cohort, where women colonized with serotype III had particularly low concentrations of serotype-specific anti-CPS IgG that did not differ from those observed in uncolonized women. The authors suggested that these low antibody levels might partly explain the over-representation of serotype III among early-onset invasive GBS isolates [39]. Our serotype distribution appears regionally distinctive. In the European DEVANI cohort, serotypes Ia, Ib, II, III and V together accounted for 88% of carriage and 94% of early-onset isolates, with serotype IV rare in both [14]. An earlier Moroccan study in Rabat found serotype IV in 5 of 67 typed maternal isolates (7%) but in 3 of 8 typed isolates from colonized newborns (38%, p=0.035), a difference the authors attributed to transmission, and which led them to describe serotype IV as emergent in that setting [18]. Our findings, almost a decade later, are consistent with that trajectory: serotype IV was one of the most frequently carried serotypes and accounted for every typed invasive isolate [17]. Serotype IV is likewise an emerging invasive serotype in other regions [25]. Because most carriers without serotype-matched antibody did not develop invasive disease, low antibody is better read as increasing susceptibility than as a sufficient cause.

### Placental transfer

Cord blood antibody closely mirrored maternal antibody, the two were strongly correlated for every serotype (r = 0.84-0.90), confirming that the newborn’s capsular antibody comes almost entirely from the mother [1]. But transfer across the placenta was incomplete, with ratios below 1 for every serotype (all p<0.001 against a ratio of 1.0), ranging from 0.80 for serotype IV to 0.50 for serotype Ib, so that roughly 50-80% of the maternal concentration reached cord blood. This finding is consistent with previous reports of naturally acquired anti-GBS IgG. In a US case-control study, transfer of anti-serotype III IgG averaged 76% and increased with gestational age, with 56% before 34 weeks, 66% at 34-36 weeks and 77% at term [8]. Similarly, transfer ratios were 66-79% after a maternal trivalent GBS vaccine [31]. In our cohort, transfer ratio was associated with gestational age (ρ=0.21, p=0.04) but not with birthweight (p=0.54) or mode of delivery (p=0.21). The incomplete transfer of specific IgG could be due to several reasons. Anti-

CPS antibody is mostly IgG2, which crosses the placenta less efficiently than the IgG1 that is mainly elicited against protein epitopes [32,37]. Identification of anti-CPS IgG subclasses would be of value to evaluate the transfer efficiency. Also, a high maternal antibody level can saturate the placental transport system and lower the transfer of antibodies [2,33]. In support of this, for instance, a mother with very high anti-serotype II antibody (about 119 µg/mL) had a cord blood level at only 0.69 µg/mL, with a ratio of 0.006. A high maternal concentration therefore does not guarantee that the newborn reaches the concentration associated with risk reduction.

### Invasive disease reflects the joint failure of two maternal protective parameters

In our previous study of the inflammatory profile in the parent cohort, we found that GBS-colonized mothers with low levels of IL-17A had an increased risk of transmitting bacteria to their neonates [20]. Combining IL-17A with serotype-matched anti-CPS IgG into a single composite separated the transmission groups (Kruskal-Wallis p=0.014): non-transmitting mothers scored highest (+2.87), mothers of symptomatic infants intermediate (-0.26), and transmitting mothers lowest (-1.66), with only the non-transmitting versus transmitting contrast reaching significance on Dunn’s test. Transmitting mothers thus fell at the low end of both parameters rather than at the point where either alone failed. The two parameters are mechanistically complementary: anti-CPS IgG mediates opsonophagocytic and complement-dependent killing of the organism, whereas the Th17/IL-17A axis drives neutrophil recruitment and mucosal clearance [34].

### Implications for maternal immunization

The link between low maternal antibody and infant disease has been known for decades [6]. A US case-control study reported that maternal anti-III IgG was globally low, with about one in five colonized mothers below 2 µg/mL and mothers of infected cases lower than mothers of healthy infants [8]. For serotype III, cord blood IgG concentration of about 0.12-0.27 µg/mL has been associated with a 75-90% reduction in the risk of invasive GBS disease, and a pooled concentration of about 0.22 µg/mL across all six serotypes was associated with an approximately 80% reduction in risk [10–13]. Published thresholds vary with sample source, assay and population, and cannot be compared directly across studies; we therefore applied the serotype-specific, onset-specific thresholds derived from the CDC case-control study throughout [13]. On that basis, only 13.0% of newborns (10/77) reached the early-onset threshold and 44.2% (34/77) the late-onset threshold, and all five invasive cases occurred in newborns below the relevant threshold. This shortfall occurred from low maternal antibody rather than from poor placental transfer. These proportions also rest on a single cord measurement at delivery; because anti-CPS IgG declines over the first three months of life [36], the proportion of infants above the late-onset threshold at the ages when late-onset disease actually occurs will be lower than the 44.2% observed at birth. Taken together, these findings support maternal immunization as a strategy in this setting, particularly where intrapartum antibiotic prophylaxis is difficult to deliver and cannot prevent late-onset disease [5, 12, 30, 31, 35]. This is particularly relevant for serotype IV, which was among the most frequently carried serotypes here and accounted for every typed invasive isolate: it is covered by the hexavalent GBS6 candidate (Ia, Ib, II, III, IV and V), the most advanced vaccine in clinical development, whose coverage therefore appears well matched to the serotypes circulating in this population.

### Strengths and limitations

This study is the first conducted in North Africa to use a paired mother-newborn design with a standardised six-plex assay and assessment in relation to GBS colonising serotype. Several limitations should be considered when interpreting the findings. Antibody concentrations were measured at a single time point, at delivery, and therefore do not capture changes in antibody levels throughout pregnancy [29]. The invasive group was small, comprising five newborns, and only three invasive isolates were available for serotyping [17]. Findings from this group should therefore be considered exploratory and require confirmation in larger North African cohorts. The transfer analysis was restricted to mother-newborn dyads with quantifiable maternal antibody concentrations, reflecting the low concentrations of naturally acquired antibody in this cohort. Consequently, placental transfer at very low maternal antibody concentrations could not be reliably assessed. In addition, the assay measures antibody binding but not functional activity; the absence of an opsonophagocytic assay therefore limits assessment of the functional capacity of the measured antibodies [27].

The risk-reduction thresholds applied in this study were derived from other populations [13], and their applicability to this North African population remains uncertain. Furthermore, although the GASTON assay was developed and validated for use with serum, it was applied to plasma samples in this study. No formal relationship between serum and plasma measurements using the GASTON MIA has been established, and this should be taken into account when interpreting the findings. This is particularly relevant to the interpretation of absolute antibody concentrations and the application of published risk-reduction thresholds, which were derived from serum measurements [13]. Classification of cord antibody concentrations above or below these thresholds should therefore be considered indicative rather than definitive. However, as the same sample matrix was used throughout the study, the use of plasma is less likely to affect internal comparisons, including between-group comparisons, maternal–cord correlations and transfer ratios. Finally, the six-plex assay did not include serotypes VI (n=2) or VII (n=6), and antibody responses associated with non-typeable strains (n=2) could not be assessed in a serotype-specific manner.

## Conclusion

In this first paired mother-newborn study of naturally acquired anti-GBS capsular immunity in North Africa, protection at birth was constrained at three sequential points: low maternal serotype-specific IgG, incomplete placental transfer for every serotype, and consequently most newborns falling beneath serotype-and onset-specific risk-reduction thresholds, including for the colonizing serotype. Passive immunity acquired through natural exposure is thus insufficient to cover the window of vulnerability in this population, providing a mechanistic rationale for maternal immunization. Two observations further inform vaccine strategy. Serotype IV was among the most frequently carried serotypes and accounted for every capsular-typed invasive isolate, a distribution more prominent than earlier European surveillance would predict and directly relevant to vaccine coverage. In parallel, a composite of maternal anti-CPS IgG and IL-17A discriminated transmission outcomes, indicating that humoral and mucosal effector mechanisms contribute jointly to neonatal risk; this analysis is exploratory, constrained by a small number of non-transmitting and invasive cases, and requires replication. Validation in larger North African cohorts, coupled with sustained serotype surveillance, will be essential to align maternal GBS vaccine composition with regional epidemiology.

## Acknowledgements

We are grateful to Raphael Simon and Natalie Silmon de Monerri (Pfizer Inc) for recommending us to perform the GASTON multiplex immunoassay work with the collaboration of Kirsty Le Doare’s team at City St George’s, University of London. This work was carried out as part of a PRD project funded by the Académie de recherche et d’enseignement supérieur (ARES). N.A, A.C and H.B.T are PhD fellows from ARES.

## Contributors

N. A. conducted the analysis and wrote the original draft. T.H and E.B performed the laboratory work, A.C. and H.B.T. assisted with the analysis. K.L. supervised the study and reviewed the manuscript. S.R. coordinated the project and supervised the study. M.C. designed the project, supervised the study, reviewed, and edited the final version. All authors contributed to the manuscript content.

